# Novel Tau Interaction Partners as Potential Biomarkers and Therapeutic Targets in Neurodegenerative Diseases

**DOI:** 10.64898/2026.11.04.764196

**Authors:** Karoline Pichlerova, Jakub Sinsky, Stefana Njemoga, Rostislav Skrabana, Adam Polak, Matej Kotasek, Petra Majerova, Martin Cente, Katarina Matyasova, Jozef Hanes

## Abstract

Alzheimer’s disease is a progressive neurodegenerative disorder leading to cognitive decline and ultimately death. One of the main hallmarks of this disease is the accumulation of pathological tau protein into neurofibrillary tangles. To better understand these pathological mechanisms, it is essential to characterize the tau protein interactome in health and disease. In this study, we performed a yeast two-hybrid screening using a human brain cDNA library, from which we identified 15 novel tau-interacting proteins that bind to full-length human tau, or its fragments: the N-terminus, microtubule-binding domain, or C-terminus. We subsequently validated these novel interactions using in vitro co-immunoprecipitation and colocalization assays with both physiological and pathological tau isoforms. Furthermore, their interactions with the human tau protein were analyzed using surface plasmon resonance. Finally, an *in silico* interactome analysis of these newly discovered tau-binding partners revealed several additional interactors that can link them to downstream processes critical to neurodegenerative pathways. Our findings offer fresh insights into tau biology and highlight novel targets for therapeutic intervention and biomarker development.

## Introduction

One of the main hallmarks of neurodegenerative diseases, specifically tauopathies including Alzheimer’s disease (AD), is the formation of neurofibrillary tangles composed of pathologically modified tau protein. This pathological environment leads to neuronal loss and cerebral atrophy, resulting in cognitive decline and, ultimately, death. AD and related tauopathies remain incurable and place an immense burden on both affected individuals and their caregivers. Globally, the number of patients suffering from AD is steadily increasing and is predicted to triple by 2050. Although several therapeutic approaches have been developed, early diagnostic markers and fully effective treatments are still lacking [1].

To achieve successful treatment with effective therapeutic agents, it is essential to identify specific biomarkers in the early stages of disease to unravel the pathological processes and mechanisms occurring before the onset of symptoms. This underscores the urgent need for a better understanding of the onset and progression of tau pathology itself.

Besides neurofibrillary tangles, another hallmark of AD is the presence of amyloid plaques composed of amyloid-beta (Aβ) peptides. Aβ are derived from the amyloid precursor protein, which physiologically serves several distinct functions. Under physiological conditions, soluble Aβ is efficiently cleared from the brain. While Aβ monomers are initially soluble, pathological accumulation triggers their misfolding. They aggregate from soluble oligomers into protofibrils, and finally into mature, insoluble fibrils that deposit as amyloid. Although senile plaques and neurofibrillary tangles represent core pathological hallmarks of AD, tau pathology is known to correlate more closely with clinical disease progression than amyloid-beta pathology [2].

Tau protein belongs to the family of microtubule-associated proteins (MAPs) [3, 4] and plays several key roles, primarily within the central nervous system. The primary physiological function of the tau protein is to bind to and stabilize microtubules. Tau protein also regulates axonal transport and synaptic plasticity. Furthermore, the role of tau protein within the neuronal nucleus includes protective interactions with DNA and RNA, thereby ensuring genomic DNA integrity [5]. At the molecular level, under pathological conditions such as AD, the tau protein becomes abnormally hyperphosphorylated and truncated. This leads to self-aggregation, which disrupts intracellular transport and protein homeostasis, ultimately resulting in neuronal death and chronic neuroinflammation [6, 7]. Although AD is rarely defined by tau pathology alone - being classified in most cases as a mixed proteinopathy [8] - the progression of tau pathology correlates strongly with cognitive decline [9]. Moreover, both the upstream events that initiate tau pathology and the downstream, tau-driven pathological processes remain incompletely understood [8]. It is highly probable that these downstream, tau-driven pathological changes within the cerebral milieu become self-sustaining, independent of the initial triggering events. Consequently, for therapeutic purposes, it is critical to identify and target these downstream pathological processes [10].

Since protein complexes and underlying protein-protein interactions (PPIs) are fundamental pillars of molecular pathways and biological processes, disruptions in the physiological tau interactome likely drive the development of pathological interactions that promote aberrant conditions and abnormal tau conformations. The relevance of this topic is underscored by the number of known tau-interacting proteins (TiPs), which have more than doubled over the last five years (n > 1100) [11]. However, most of them have been identified through high-throughput screening methods and have not been validated by targeted experiments. Moreover, to fully comprehend the implications of these interactions in pathogenesis, it is necessary to determine whether they contribute to neuroprotection or rather facilitate neurodegeneration. Thorough characterization of tau interactions under both physiological and pathological conditions is essential for understanding the processes underlying primary tauopathies and for identifying potential biomarkers.

To date, several approaches have been applied to discover novel tau interaction partners. However, most of them were performed as large proteomic screenings and did not proceed to validation [12, 13]. In some cases, post-screening studies focused exclusively on a single downstream protein [14, 15].

Recently, we identified 175 potential novel and known tau-interacting proteins in transgenic and wild-type rat brains using an *in vivo* crosslinking approach coupled with mass spectrometry. In subsequent validation experiments, three candidates were confirmed as novel tau-interacting partners. Based on their known functions and involvement in neurodegenerative or psychiatric disorders, we proposed a role for these proteins in tau pathology [16].

To expand on our previous findings and identify additional TiPs, we employed a complementary approach: we screened a human brain cDNA library against the longest human tau isoform and its distinct fragments as baits by the yeast two-hybrid system (Y2H). Identified novel TiPs were used for comprehensive bioinformatic analysis and further validated by biochemical methods.

## Material and methods

### Chemicals

Basic reagents and chemicals were purchased from Merck unless otherwise stated. Restriction enzymes, Phusion DNA polymerase and T7-ligase enzymes were purchased from New England Biolabs (NEB; England). L-tryptophan, L-lysine, and L-leucine were purchased from Sunrise Science Products (USA). L-threonine, L-phenylalanine, L-valine, and L-isoleucine were purchased from Alfa Aesar (USA). Yeast Nitrogen Base and agar were purchased from Formedium (England). The Aureobasidin A, X-α-gal, and normalized human brain cDNA library were purchased from Takara Bio (Japan). The Nonidet P40 Substitute (NP-40) was purchased from Honeywell Fluka™ (USA), cOmplete protease inhibitor cocktail tabs from Roche (Germany), Fluoroshield™ with DAPI mounting medium and SuperSignal™ West Pico PLUS Chemiluminescent Substrate were purchased from ThermoFisher Scientific (USA). Full-length human tau protein isoform 2N4R was produced at the Institute of Neuroimmunology [17]. The backbone plasmid for the yeast two-hybrid system (pGBKT7, Clontech) was obtained from Takara Bio. For protein expression in Chinese hamster ovary (CHO) cells, the pCMV-3UTR plasmid was used [18].

### Antibodies

Mouse anti-tau monoclonal antibodies (mAb, recognizing epitopes of the longest isoform of human tau isoform 2N4Rtau): DC31N (aa 1-30), DC25 (aa 347-353), DC18 (aa 168-181, specific for human tau), DC190 (aa 368-376) and mouse anti-tau polyclonal antibody (pAb) recognizing α-tubulin were obtained from AXON Neuroscience SE, Slovakia. Other antibodies used in the study: rabbit anti-ATP1B3 pAb (11142-1-AP), rabbit anti-BPGM pAb (17173-1-AP), mouse anti-PCNA mAb (10205-2-AP), rabbit anti-PCCA pAb (21988-1-AP), rabbit anti-SGCB pAb (11936-1-AP), rabbit anti-PDE4DIP pAb (12637-1-AP), rabbit anti-ATP1B1 pAb (15192-1-AP), mouse anti-His-Tag mAb (66005-1-Ig) (all from Proteintech, Germany), rabbit anti-NCAM1 pAb (MA5-33055), rabbit anti-NGLY1 pAb (PA5-110030), mouse mAb-CACYBP1 mAb (68161-1-Ig), rabbit anti-BLTP3B pAb (PA5-115467), rabbit anti-IL1RAcP pAb (PA5-76791) (all from Invitrogen, Thermofisher Scientific, USA). Secondary antibodies used in our experiments were goat anti-rabbit pAb and goat anti-mouse pAb, both conjugated to horseradish peroxidase (Agilent, USA), goat anti-mouse pAb and goat anti-rabbit pAb conjugated with Alexa Fluor® 488 or Alexa Fluor® 546, respectively (ThermoFisher Scientific, USA).

### Biological material

Human Alzheimer’s brain tissue (female, neuropathology: Alzheimeŕs disease, Braak stage VI) and healthy brain tissue (female, no psychiatric symptoms, no dementia, Braak stage I) were obtained from the Queen Square Brain Bank for Neurological Disorders (London, UK). All samples were obtained with informed patient consent and with approval from the local ethical committees.

Animals were housed under standard laboratory conditions with free access to water and food and maintained under diurnal lighting. All experiments on animals were carried out according to the institutional animal care guidelines conforming to international standards (Directive 2010/63/EU) and were approved by the State Veterinary and Food Committee of Slovak Republic (RO-6303-4/2025-220/4504/2025). Humane endpoints based on body weight loss and body condition scoring were in place. No unexplained mortality occurred in these studies. All animal experiments were monitored and approved by the Institutional Ethics Committee (Ethics Committee of the Institute of Neuroimmunology, Slovak Academy of Sciences). We used 5-month-old transgenic rats expressing human truncated tau protein (aa151-391/4R) [19] and age-matched wild-type rats with the same SHR genetic background.

Monocytes for cDNA library preparation were obtained from whole blood of two voluntaries, under the ethical commissions (Etická komisia Univerzitnej nemocnice Bratislava UNB, Nemocnica sv. Cyrila a Metoda) approval EK4/4/2026.

### Strains

*Saccharomyces cerevisiae* strains used in the screening was the Y2H Gold strain (genotype: MATa, trp1-901, leu2-3,112, ura3-52, his3-200, gal4Δ, gal80Δ, LYS2::GAL1UAS–Gal1 TATA–His3, GAL2UAS–Gal2TATA–Ade2URA3::MEL1 UAS–Mel1 TATA AUR1-C MEL1) containing the bait protein and Y187 strain (genotype: MATα, ura3-52, his3-200, ade2-101, trp1-901, leu2-3, 112, gal4Δ, gal80Δ, met–, URA3::GAL1UAS–Gal1TATA–LacZ, MEL1) carrying the library. The XL1-Blue *Escherichia coli* strain was used as a bacterial carrier of cloned recombinant plasmids (genotype: recA1 endA1 gyrA96 thi-1 hsdR17 supE44 relA1 lac [F proAB lacIqZΔM15 Tn10 (Tetr)]). *S. cerevisiae* stains were obtained from Takara Bio (Y187-630457; Y2H Gold - 630498) and XL1-Blue *E. coli* were obtained from Agilent (200249).

### Yeast two-hybrid screening

The normalized human brain cDNA library was acquired commercially from TakaraBio (Japan) with a size of 3.56 × 10^6^ genes and a library titer of 8 × 10^7^ cfu/mL clones. The screening was carried out according to the manufacturer’s protocol, with the following modifications. Synthetic defined (SD) media for the yeast two-hybrid system were prepared according to the Cold Spring Harbor Protocols [20]. The SD dropouts media without tryptophan (SD-W), without tryptophan and leucine (SD -L, W; double drop-out; DDO) and without tryptophan, leucine, adenine, histidine (SD -L, W, A, H; quadruple drop-out; QDO) were used to select transformants positive for bait, for bait and prey, and for interaction of bait and prey, respectively. The transformation of the bait plasmid (pGBKT7) was carried out using the chemical TE/LiAc protocol [21] and plated on bait-selective media (SD–W). The transformants were allowed to grow for 3-4 days at 30°C. The mating procedure was carried out for 20 h. The cell suspension was plated on fifty 150 mm plates with QDO medium in a volume of 100-200 μl per plate and incubated for 5 days at 30°C. The grown colonies were patched to more stringent plates (QDO supplemented with Aureobasidin A and X-α-gal) and grown at 30°C for another 5 days. Subsequently, blue colonies were patched twice onto double-dropout (DDO) plates with X-α-gal to visualize plasmid segregation. The colonies containing the segregated target-prey plasmid were grown for 2 days in 3 mL of plasmid-selective medium. The cells were resuspended in 200 μl of lysis buffer (200 mM NaOH, 1% SDS (w/v)) and were additionally disrupted by 3 rounds of vortexing with 200 μl of glass beads (500–600 μm) for 1 minute, followed by chilling on ice for 5 minutes. To remove duplicates, PCR was performed on prey plasmids, and the resulting inserts of the same size were subjected to restriction analysis with the BfaI enzyme. Inserts with the same cleavage pattern were removed, and unique plasmids were transformed into *E. coli* and plated onto Ampicillin plates to rescue the prey plasmids. Prey plasmids were isolated and co-transformed with the bait pGBKT7 constructs with tau proteins, including empty pGBKT7 as a negative control, into the *S. cerevisiae* Y2H Gold strain using the TE/LiAc transformation protocol. After growth on DDO medium, two single colonies were picked and patched onto DDO supplemented with X-α-gal and QDO with Aureobasidin A and X-α-gal (QDO/A/X). The plates were grown at 30°C for up to 5 days. Their growth and color (on QDO) and color changes (on DDO) were monitored and compared. The prey plasmids from true-positive clones (with growth and color changes on QDO/A/X) were sequenced.

### Co-immunoprecipitation

The whole procedure, including protein extraction, sample handling, and co-immunoprecipitation (co-IP), was performed at 4°C. Brainstem tissues from control and transgenic rats [19], AD and healthy human cortices were homogenized in a mechanical homogenizer in TNL buffer (50 mM Tris pH 7.4, 150 mM NaCl, 1mM EDTA, 1% NP-40, supplemented with protease inhibitor cocktail, Complete EDTA-free (Roche, Germany)), with a ratio of 3 mL of TNL buffer for 0.5 g of tissue. The homogenates were incubated by head-over-head rotation for 30 min, then centrifuged at 15,000 × g for 20 min, and the supernatants were collected and stored on ice. The extraction process was repeated once more, and the supernatants were pooled.

Protein extracts were then filtered through a 0.22 µm filter and diluted twice with WBNP0.1 buffer (50 mM Tris pH 7.4, 150 mM NaCl, 0.1% NP-40). The co-IP was carried out as described previously with some modifications [16]. Briefly, a mixture of three anti-tau mAbs, DC18, DC25, and DC190, immobilized on CNBr Sepharose (Cytiva, Sweden), was used to capture tau complexes. Eluates from the columns were precipitated with 10% trichloroacetic acid for 1 h on ice. After precipitation, samples were centrifuged at 20,000 × g for 15 min at 4°C, washed twice with ice-cold 96% ethanol, and dried at room temperature for 30 min. The dried samples were dissolved in SDS protein loading buffer and analyzed by Western blotting. Dilutions of the antibodies in experiments were as follow - ATP1B1 1:1000; ATP1B3 1:1000; BLTP3B 1:1000; BPGM 1:1000; CACYBP 1:12000; IL1RAP 1:1000; NCAM1 1:1000; NGLY1 1:1000; PCCA 1:1000; PCNA 1:5000; PDE4DIP 1:1000; SGCB 1:1000.

### Colocalization experiments

For all experiments, three types of SH-SY5Y human neuroblastoma cell lines were used: 1) parental SH-SY5Y, 2) SH-SY5Y with inducible expression of tau151-391/4R [22], and 3) SH-SY5Y with inducible expression of 2N4R-Tau [22]. The cells were maintained in 1:1 mix of Minimum essential medium and Ham’s F12 medium (PAA laboratories GmbH, Austria) supplemented with 10% fetal bovine serum (GIBCO-Invitrogen, USA), 2 mM L-glutamine (PAA laboratories GmbH, Austria), 1% non-essential AA (PAA laboratories GmbH, Austria), 50 µg/mL gentamycin (PAA laboratories GmbH, Austria), 50 µg/mL G418 (GIBCO-Invitrogen, USA), 100 µg/mL HygromycinB (Invitrogen, USA) and 10 ng/mL doxycycline (Sigma-Aldrich, USA). Cell cultivation was performed at 37°C in a 5% CO_2_ atmosphere. Tau protein expression was induced by culturing cells in a doxycycline-free medium for 3 days before use in experiments. After 3-day cultivation without doxycycline, SH-SY5Y cell cultures were transferred to 8-well Lab-Tek™ II Chamber Slides™ (ThermoFisher Scientific, USA), rinsed briefly with phosphate-buffered saline (PBS) and fixed in 4% paraformaldehyde in PBS for 10 min at 37°C. After rinsing 3 times with PBS, cells were permeabilized for 2 min with 0.3% Triton X-100 in PBS, washed, and blocked for 1 h at 25°C in blocking buffer (0.1% Triton X-100 and 5% bovine serum albumin in PBS). Fixed cells were incubated overnight with primary antibodies diluted in the blocking buffer at 4°C. The next day, cells were rinsed 3 times with PBS and incubated with the corresponding fluorescently labeled secondary antibodies. Stained preparations were mounted with Fluoroshield DAPI mounting medium (Sigma-Aldrich, USA) and examined under an LSM 710 confocal microscope (Zeiss, Germany) at 63 × magnification using 488 nm and 561 nm lasers to excite fluorophores. Colocalization was detected by pixel-to-pixel overlay of both images containing green and red emission signals. Dilutions of the antibodies in the colocalization experiments were as follows: ATP1B1 1:100; ATP1B3 1:500; BLTP3B 1:100; BPGM 1:200; CACYBP 1:400; IL1RAP 1:200; NCAM1 1:200; NGLY1 1:200; PCCA 1:200; PCNA 1:200; PDE4DIP 1:200; SGCB 1:500.

### Molecular cloning of target genes

Peripheral blood mononuclear cells were isolated using MACSprep PBMC Isolation Kit (Miltenyi Biotec, USA, Cat# 130-139-976) and monocytes by Pan Monocyte Isolation Kit (Miltenyi Biotec, USA, Cat# 130-096-537). Total RNA was isolated using the RNeasy Mini Kit according to the manufacturer’s recommendations (Qiagen, Germany, Cat# 74104). First-strand cDNA synthesis was performed using the High-Capacity cDNA Reverse Transcription Kit (Applied Biosystems, USA, Cat# 4368813) following the manufacturer’s protocol. Genes of selected proteins were obtained by PCR amplification with specific primers (supplementary table 1) using Phusion^®^ DNA Polymerase (New England Biolabs). PCR products were cloned into an expression vector pCMV-3UTR cut with AgeI and XhoI restriction endonucleases [18]. All constructs were verified by DNA sequencing.

### Protein expression in CHO cells

ExpiCHO-S cells (Gibco, USA) were cultivated in the BalanCD CHO Growth A Medium (Fujifilm Biosciences, Japan) supplemented with 6 mM GlutaMAX (ThermoFisher Scientific, USA) in a suspension culture. The cells were grown in the presence of antibiotics (0.06 mg/mL penicillin and 0.1 mg/mL streptomycin, Sigma-Aldrich, St. Louis, MO, USA) at 37°C and 8% CO_2_ in a humidified atmosphere with constant shaking at 130 rpm on a 19 mm orbital shaker (Advanced Dura-Shaker, VWR Avantor, PA, USA). Non-baffled polycarbonate Erlenmeyer flasks with vent cap (Corning, Corning, NY, USA) or SIMAX glass Erlenmeyer flasks (Kavalierglass, Praha, Czech Republic) with DURAN vented screw cap (DWK Life Sciences, Wertheim, Germany) were used for cultivation. The cells were regularly passaged after reaching a density of 5 × 10^6^ per mL of culture.

Transfection with expression vectors containing target genes was accomplished using polyethylenimine (PEI) gene delivery. 24 hours before transfection, growth medium was replaced with BalanCD CHO Transfectory medium (Fujifilm Biosciences), supplemented with 8 mM GlutaMAX, and cells were seeded at a density of 2.5 × 10^6^ cells/mL. Just prior to transfection, the culture density was adjusted to 5 × 10^6^ cells/mL, and the transfection mix of DNA and PEI in transfectory medium was added as described previously [18]. The transfected cells were incubated at 37°C with shaking as above. On the day following transfection, the culture was supplemented with 15% (v/v) BalanCD CHO Feed 4 (Fujifilm Biosciences, Japan), penicillin and streptomycin to final concentrations of 0.06 mg/mL and 0.1 mg/mL, respectively, and sodium butyrate to a final concentration of 0.9 mM. The culture was then transferred to a 32°C incubator with 5% CO₂, and cultivation was continued for 5–7 days. The culture media were collected and stored at -20°C.

### Protein purification

Tau-interacting partners were expressed as His-tagged proteins in the CHO expression system with secretion into the culture medium [23]. The whole purification procedure was carried out at 4°C. The harvested media were adjusted to pH 7.0, loaded onto a 5 mL HisTrap™ column (Cytiva, USA), and washed with a buffer containing 20 mM Tris (pH 8.0) and 500 mM NaCl. Bound proteins were eluted using the same buffer supplemented with 250 mM imidazole. Collected fractions were analyzed by 12% SDS-PAGE. The concentrations of the target proteins were determined by Western blotting with an anti-His antibody (1:10000), using His-tagged rat Angulin 1 as the protein standard, whose concentration was estimated by BCA assay and confirmed by Coomassie blue staining of the SDS-PAGE gel.

### Surface plasmon resonance

Interaction kinetics of tau binding partners were measured at 25°C in PBS-P+ buffer (PBS containing 0.05% Tween-20) by a Biacore 1K+ system running Biacore Insight Control software (version 7.0.9.1992; Cytiva, Uppsala, Sweden). 2N4R-Tau was immobilized on a Series S sensor chip CM5 (Cytiva) by amine coupling. A single-cycle kinetics protocol was employed to measure the binding kinetics of multiple identified binding partners, including BPGM, IL1RAP, SNAPIN, and PITHD1. The binding cycles consisted of four subsequent injections of tau-binding partners, each followed by a 120-second contact time, followed by a final 180-second dissociation. The flow rate was set to 100 µl/min. Regeneration of the chip surface was accomplished by three six-second-long injections of 100 mM HCl. Specific monoclonal antibodies with well-characterized tau epitopes were used as controls to confirm that 2N4R-Tau remained accessible after immobilization to the CM5 surface. Binding kinetics were calculated in Biacore Insight. Evaluation software (version 7.0.9.1992).

### Bioinformatic analysis

Tau protein interactions were further explored through bioinformatic analysis, evaluating 15 proteins examined in this study, along with 6 tau-interacting partners identified in our previous analyses [16, 24]. PPIs for the 21 investigated proteins were retrieved from the Integrated Interactions Database (IID) version 2025-05 (https://iid25.ophid.utoronto.ca) [25], including Alzheimer’s disease annotations. The network was built, annotated, and analyzed using NaViGaTOR version 3.0.19 [26]. To identify the most relevant proteins within the network, we computed node degree, which reflects the number of PPIs among interacting partners. Highly interconnected proteins among tau interaction partners (node degree >7) were highlighted by name and node size and sorted by Gene Ontology (GO) Function. The original network was subsequently prioritized to visualize interactions associated with Alzheimer’s disease (blue edges). Figures were exported in SVG format and finalized in CorelDraw version 24.5.0.731 (Corel Corporation, Ottawa, ON, Canada).

## Results

### Yeast two-hybrid screening of a human brain cDNA library

The Y187 strain contained a human brain cDNA library, with the proteins fused to the Gal4 system’s activation domain, acting as prey in this system. The Y2HGold strain contained tau protein and tau fragments fused to the binding domain, functioning solely as baits in the system. The expression and auto-activation of tau protein and its fragments were checked in the system prior to the screenings. The proteins utilized as baits in the screening included human 2N4R tau (full-length human physiological tau, aa 1-443), N-terminus of tau (aa 2-243), microtubule binding domain of tau (aa 244-368), tau151-391 (aa151-391) and C-terminus of tau (aa 369-441). After screening, autoactivating false positives were excluded. The number of diploids screened and the mating efficiency varied across screenings; the data are presented in the supplementary data (Table S2). After this first validation step, 15 new proteins were discovered. The number of isolated, unique, and true-positive clones, as well as the number of new proteins, for each screening is shown in Table 1. The list of novel tau interaction partners, including their full names and descriptions of their known roles in neurodegeneration, is shown in Table 2. A screening with the tau151-391 fragment resulted in complete growth inhibition post-mating; therefore, this experiment was terminated.

**Table 1.** The number of clones isolated in each screening and the resulting amount of novel tau interaction partners after each screening.

| **Tau protein/Tau fragment** | **Clones isolated after screening** | **Unique clones** | **True positive clones** | **Identified interaction partners** |
|---|---|---|---|---|
| 2N4R tau<br>(aa 1-441) | 69 | 48 | 3 | 2 |
| N-terminus<br>(aa 2-243) | 65 | 63 | 6 | 3 |
| MTBD<br>(aa 244-368) | 111 | 97 | 16 | 3 |
| C-terminus<br>(aa 369-441) | 247 | 210 | 33 | 8 |
| Summarized | 492 | 418 | 58 | 15* |

**Note:** One protein was discovered in screenings against two different tau fragments (2N4R and C-terminus)

**Table 2.** Novel tau interaction partners identified in this study.

| **Identified protein from Y2H screening** | **Gene** | **Identified with** | **K** | **Function and neurodegeneration** | **Citations** |
|---|---|---|---|---|---|
| ATPase Na+/K+ Transporting Subunit Beta 1 | ATP1B1 | C-terminus, Tau40 | No | non-catalytic subunit of Na+/K+-ATPase pump; regulates its stability; dysregulation is identified in neurodegenerative omics profiles | 27-29 |
| Sodium/potassium-transporting ATPase subunit beta-3 | ATP1B3 | N-terminus | No | non-catalytic subunit of Na+/K+-ATPase pump; regulates its plasma membrane targeting; present in a specific subset of neurons | 27, 30 |
| Bridge-like lipid transfer protein family member 3B isoform a | BLTP3B | MTBR | Yes | mediates inter-organelle lipid flux at membrane contact sites; disruption impairs membrane homeostasis and synaptic signaling; mutations are linked to early-onset Parkinson’s disease | 31 |
| Bisphosphoglycerate mutase | BPGM | C-terminus | Yes | enzymatic activity; regulation of erythrocyte hemoglobin-oxygen affinity and glycolytic/serine metabolic fluxes; activity is increased in AD and dementia; potential role in neurodegeneration through disrupted neuronal glucose and glycogen metabolism | 32-36 |
| Calcyclin-binding protein isoform 1 | CACYBP | MTBR | Yes | regulatory protein forming a complex involved in proteasomal degradation and cytoskeletal dynamics; complex participates in AD in tau dephosphorylation | 37-40 |
| Interleukin-1 receptor accessory protein (isoform 2; soluble) | IL1RAP | N-terminus | Yes | blocks IL-1 signaling cascades; elevated serum levels correlate with cognitive impairment | 41-47 |
| Neural cell adhesion molecule 1 isoform 27 precursor | NCAM1 | MTBR | Yes | adhesion molecule; modulates neurite outgrowth; in AD levels decline in the cortex; its polysialylated form correlates inversely with tau load | 48-53 |
| Peptide-N(4)-(N-acetyl-beta-glucosaminyl)asparagine amidase | NGLY1 | N-terminus | Yes | enzymatic activity; participates in the endoplasmic reticulum-associated degradation pathway; dysfunction leads to mitochondrial defects, neuronal protein aggregation and tau seeding | 54-62 |
| Propionyl-CoA carboxylase alpha chain (mitochondrial) | PCCA | C-terminus | Yes | mitochondrial enzyme; linked to metabolic injury and oxidative stress; interacts directly Aβ40/42; its associated circular RNA binds miR-138-5p regulating tau phosphorylation | 63-66 |
| Proliferating cell nuclear antigen | PCNA | C-terminus | Yes | essential for post-mitotic neuronal repair; part of the DNA damage response; in AD is upregulated during aberrant tau-induced cell-cycle re-entry; accumulates in the cytoplasm prior to apoptosis | 67-74 |
| Myomegalin | PDE4DIP | C-terminus | Yes | centrosomal/Golgi-anchoring protein regulating neurogenesis and microtubule assembly; its genetic polymorphisms act as risk or protective factors in frontotemporal dementia, vascular dementia, and AD | 75-79 |
| PITH domain-containing protein 1 | PITHD1 | C-terminus | Yes | modulates trans-synaptic signaling, transcription factor regulation, and pro-inflammatory proteostatic shifts; selectively upregulated in the AD olfactory tract; potential proteomic biomarker for early-stage neurodegeneration | 80-82 |
| Beta-sarcoglycan | SGCB | C-terminus | No | links the extracellular matrix to the cytoskeleton; drives postsynaptic receptor assembly in the GABAergic pathway; disruption destabilizes the dystroglycan complex | 83 |
| SH3-containing GRB2-like protein 3-interacting protein 1 | SGIP1 | C-terminus | Yes | endocytic adaptor mediating synaptic vesicle recycling and cannabinoid signaling; present in stress granules; loss of function is associated with cognitive dysfunction and early-onset Parkinsonism | 84, 85 |
| SNARE-associated protein Snapin | SNAPIN | Tau40 | Yes | participates in synaptic vesicle release and endosomal-lysosomal trafficking; its deficiency results in axonal deficiency, exacerbating AD-associated autophagic stress | 86-92 |

Note: **K, known in neurodegeneration**

### Protein interaction network for potential tau-interacting partners

To further characterize the biological processes associated with the identified tau-interacting partners, we conducted a bioinformatic analysis. The analysis explored a panel of 21 proteins, including 15 identified by this study and 6 additional proteins that showed direct interactions with tau, as described in our previous work [16, 24]. A PPI network of the investigated proteins was analyzed to identify the most connected proteins and to reveal their associations with neurodegeneration. The resulting network comprised 4685 proteins with 6902 interactions, of which 342 were annotated as relevant to Alzheimer’s disease. In this PPI network, 4 proteins were identified as known AD-specific interactors (PCNA, NCAM1, NPTX1, and IL1RAP). Eight of these 21 queried proteins had no previously reported direct interaction with tau in the IID database (BLTP3B, BPGM, IL1RAP, NGLY1, PCCA, PDE4DIP, SGCB, and SNAPIN). Furthermore, the network analysis identified highly connected proteins that interact with both tau and the TiPs. This group of shared interacting proteins included those associated with various neurodegenerative diseases, such as APP (AD), PRKN and LRRK2 (Parkinson’s disease), PRNP (prion diseases), VCP and CCNF (Amyotrophic lateral sclerosis), and ATXN1 (Spinocerebellar ataxia), highlighting the potential role of the examined tau-interacting partners in neurocognitive disorders (Fig. 1). In summary, the analysis revealed numerous proteins involved in protein homeostasis, ubiquitin-mediated degradation, heat shock and cellular stress responses, Wnt signaling, cytoskeletal organization, neuronal signaling, cell cycle regulation, and cell survival.

**Fig. 1:**
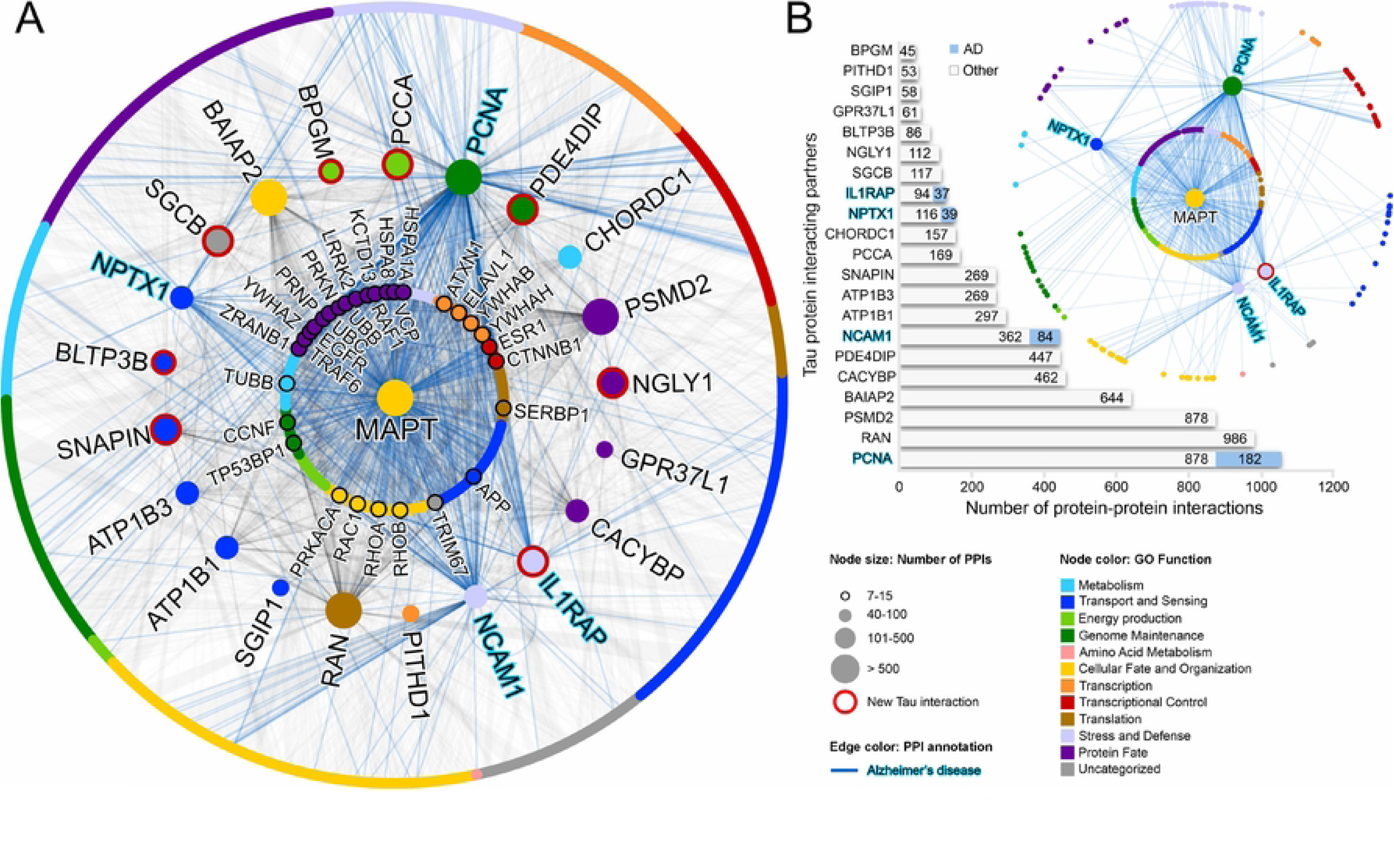
Protein-protein interaction network of tau protein-interacting partners (A). The 21 examined proteins are aligned in the middle circle and annotated with symbols. The outer circle represents proteins that interact only with the examined tau partners. The inner circle represents all proteins that interact with both tau and the 21 proteins. Node color corresponds to Gene Ontology (GO) Function, node size represents the number of interacting partners, and blue edges indicate interactions relevant to Alzheimer’s disease. Red node outline highlights a new Tau interaction partner. (B) A bar graph showing the number of interactions for each protein and the number of Alzheimer’s disease-relevant PPIs.

### Validation of novel tau interaction partners

The newly identified tau interaction partners were further validated by co-immunoprecipitation, colocalization, and, for selected proteins, by surface plasmon resonance.

**Co-immunoprecipitation** was conducted using human brain cortices from Alzheimer’s disease patient and healthy individual, and rat brainstem tissues from both spontaneous hypertensive rats (SHR), serving as controls, and transgenic rats with Alzheimer’s disease pathology (SHR72) [19]. Samples were subjected to co-IP using a mix of three pan-tau mAbs that recognize both rat and human tau proteins. The presence of newly identified TiPs in IP samples was analyzed using anti-TiP-specific antibodies.

In tau immunoprecipitates from rat brain tissues, we confirmed interactions between tau proteins and ATP1B1, NGLY1, NCAM1, BLTP3B, SGCB and IL1RAP. All proteins were present in both SHR and SHR72 brainstem samples, showing that those analyzed TiPs interact with both endogenous rat tau and human 151-391/4R tau proteins. The IL1RAP antibody reacted with an additional band at around 100 kDa, which appears to be the membrane isoform of this protein. Proteins ATP1B3, BPGM, CACYBP, PCCA, PCNA, and PDE4DIP were not detected in any of the immunoprecipitates (PDE4DIP showed a positive band at 50 kDa, significantly smaller than the expected 265 kDa) (Fig. 2A).

**Fig. 2:**
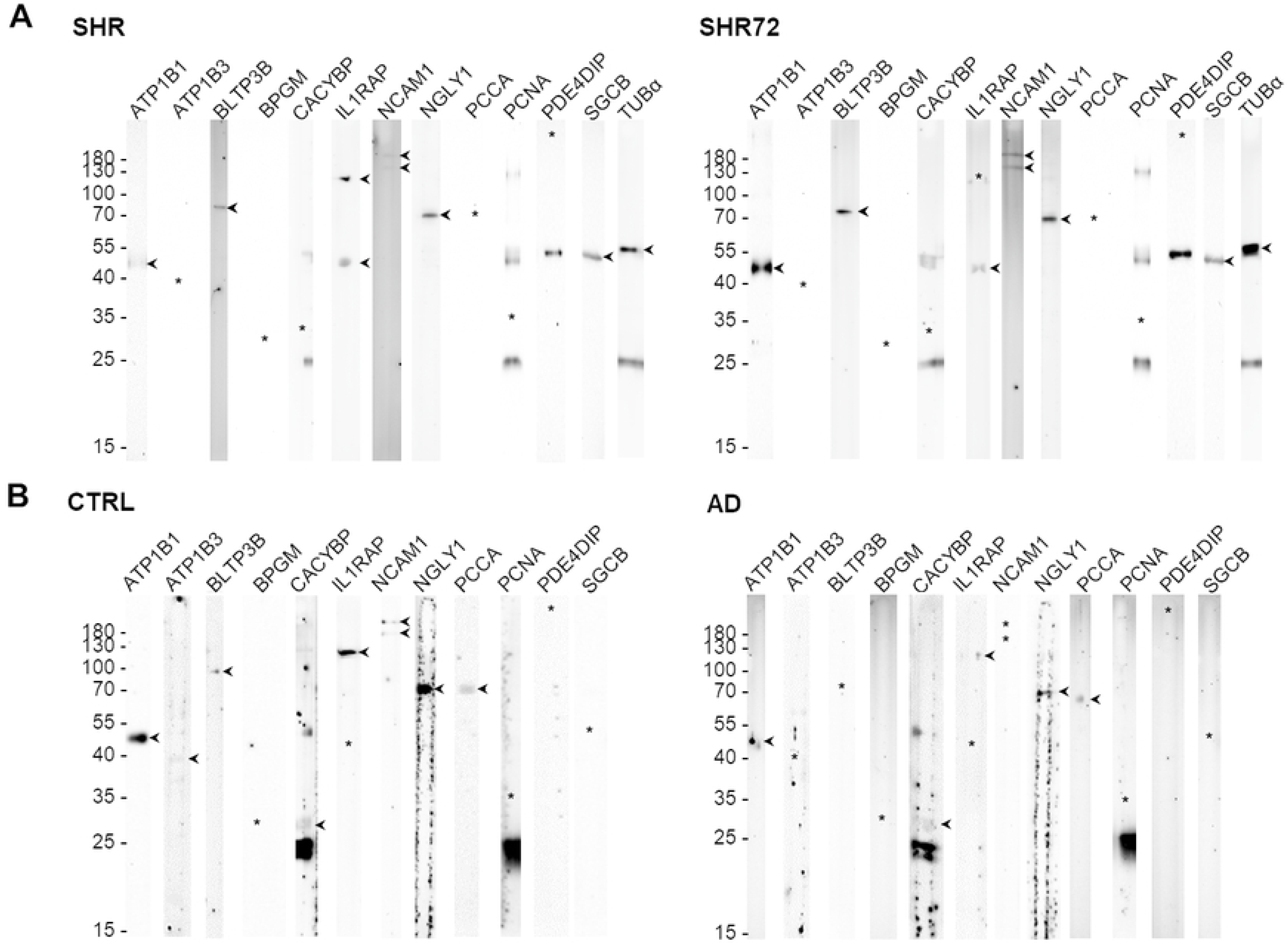
Validation of novel tau interaction partners by co-immunoprecipitation. A) Results from co-IP from rat SHR and SHR72 brainstem tissues. B) Results from co-IP from healthy human and AD tissues. Positive results are marked with arrows, and missing bands are shown as asterisks.

Co-IP from the human samples confirmed the presence of ATP1B1, CACYBP, NGLY1, IL1RAP and PCCA in both healthy and AD donors. ATP1B3, BLTP3B and NCAM1 proteins were identified only in healthy control samples. In contrast to rat samples, we identified only the membrane form of IL1RAP in both human samples, whereas the soluble, low-molecular-weight isoform of this protein was absent. In the healthy brain, we observed a triple band for ATP1B3 around its expected molecular weight of 40-55 kDa (Fig. 2B).

We confirmed the presence of ATP1B1 and NGLY1 across all co-IP samples from both rat and human. Additionally, we confirmed the presence of BLTP3B and NCAM1 in both SHR and human control samples. In staining with antibodies against CACYBP and PCNA, two additional bands (25 kDa and 50 kDa) were observed, resulting from non-specific binding of secondary anti-mouse antibody.

#### Colocalization experiments

We analyzed the interaction of tau proteins with newly identified TiPs in three SH-SY5Y cell lines: Control cell line (parental SH-SY5Y, expressing human endogenous tau protein at low concentration), cell line overexpressing human 2N4R tau (SH-SY5Y-2N4RTau) [22], and cell line overexpressing the truncated pathological form of human tau protein (SH-SY5Y-ADTau) [22]. We found that ATP1B1, BLTP3B, BPGM, PCCA, and SGCB proteins colocalized with both human 2N4R tau and 151-391tau proteins. Furthermore, we identified that NGLY1, PCNA, and PDE4DIP interacted only with human truncated tau 151-391tau (Fig. 3), and NCAM1 colocalized only with human 2N4R tau (Fig. S1). For the three tested proteins, ATP1B3, CACYBP and IL1RAP, we could not confirm any interaction with tau proteins in colocalization experiments (Fig. S2). We did not observe any colocalizations of the tested novel TiPs in the parental SH-SY5Y cell line, except for ATP1B1, which showed a weak signal (Fig. S3). The validation results for the selected proteins, using co-immunoprecipitation and colocalization, are summarized in Table 3.

**Fig. 3:**
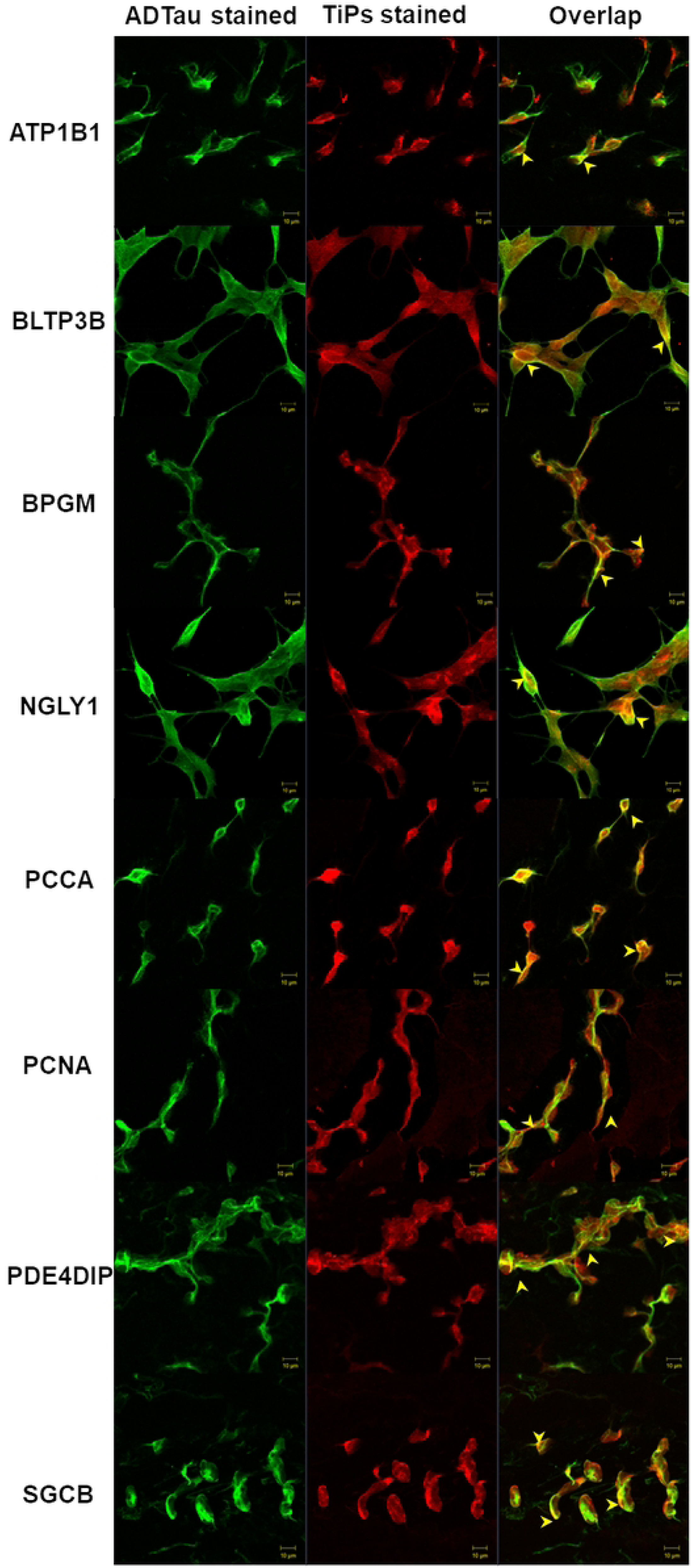
Colocalizations of potential TiPs with human 151-391tau in SHSY-5Y-ADtau cell line. The proteins ATP1B1, BLTP3B, BPGM, NGLY1, PCCA, PCNA, PDE4DIP and SGCB colocalized with tau proteins expressed in SHSY-5Y-ADtau cell line. Cells were fluorescently stained with anti-tau (green) and anti-investigated candidate (red) antibodies. Colocalization of tau with its partner is seen in yellow (selected regions highlighted by yellow arrows) as a result of green and red emission signal overlay. A ten-micrometer scale bar is indicated.

**Table 3.** Summary of *in vitro* validation experiments.

| **Proteins** | **Gene** | **Co-immunoprecipitation** |  |  |  | **Colocalization** |  |  |
|---|---|---|---|---|---|---|---|---|
|  |  | **SHR** | **SHR72** | **human** | **AD** | **wt** | **2N4R** | **ADTau** |
| ATPase Na+/K+ Transporting Subunit Beta 1 | ATP1B1 | + | + | + | + | weak | + | weak |
| Sodium/potassium-transporting ATPase subunit beta-3 | ATP1B3 | - | - | + | - | - | - | - |
| bridge-like lipid transfer protein family member 3B isoform a | BLTP3B | + | + | + | - | - | + | + |
| Bisphosphoglycerate mutase | BPGM | - | - | - | - | - | weak | weak |
| calcyclin-binding protein isoform 1 | CACYBP | - | - | + | + | - | - | - |
| Interleukin-1 receptor accessory protein (isoform 2) | IL1RAP | + | + | - | - | - | - | - |
| Peptide-N(4)-(N-acetyl-beta-glucosaminyl)asparagine amidase | NGLY1 | + | + | + | + | - | - | + |
| neural cell adhesion molecule 1 isoform 27 precursor | NCAM1 | + | + | + | - | - | weak | - |
| Propionyl-CoA carboxylase alpha chain (mitochondrial) | PCCA | - | - | + | + | - | + | + |
| Proliferating cell nuclear antigen | PCNA | - | - | - | - | - | - | weak |
| Myomegalin | PDE4DIP | - | - | - | - | - | - | + |
| Beta-sarcoglycan | SGCB | + | + | - | - | - | + | + |

**Notes:** +, interaction identified; weak, weak signal; -, interaction not confirmed

#### Surface plasmon resonance interaction analysis of TiPs with human 2N4R tau

Five newly identified TiPs: PCNA, BPGM, IL1RAP, PITHD1 and SNAPIN were analyzed by SPR to assess their direct binding to tau. As a negative control, rat Angulin 1 was used. Interaction analysis was performed using purified recombinant His-tagged proteins expressed in CHO cells. SPR analyses demonstrated that full-length tau protein (2N4R tau) interacted with BPGM, ILRAP1, PITHD1 and SNAPIN, whereas no interaction was detected with PCNA and a negative control, Angulin 1 (Fig. 4). Kinetic parameters from quantitative SPR assessments, presented in Table 4, showed that the lowest equilibrium dissociation constant (KD) was observed for the full-length tau-SNAPIN interaction, indicating that this complex exhibited the highest binding affinity.

**Fig. 4:**
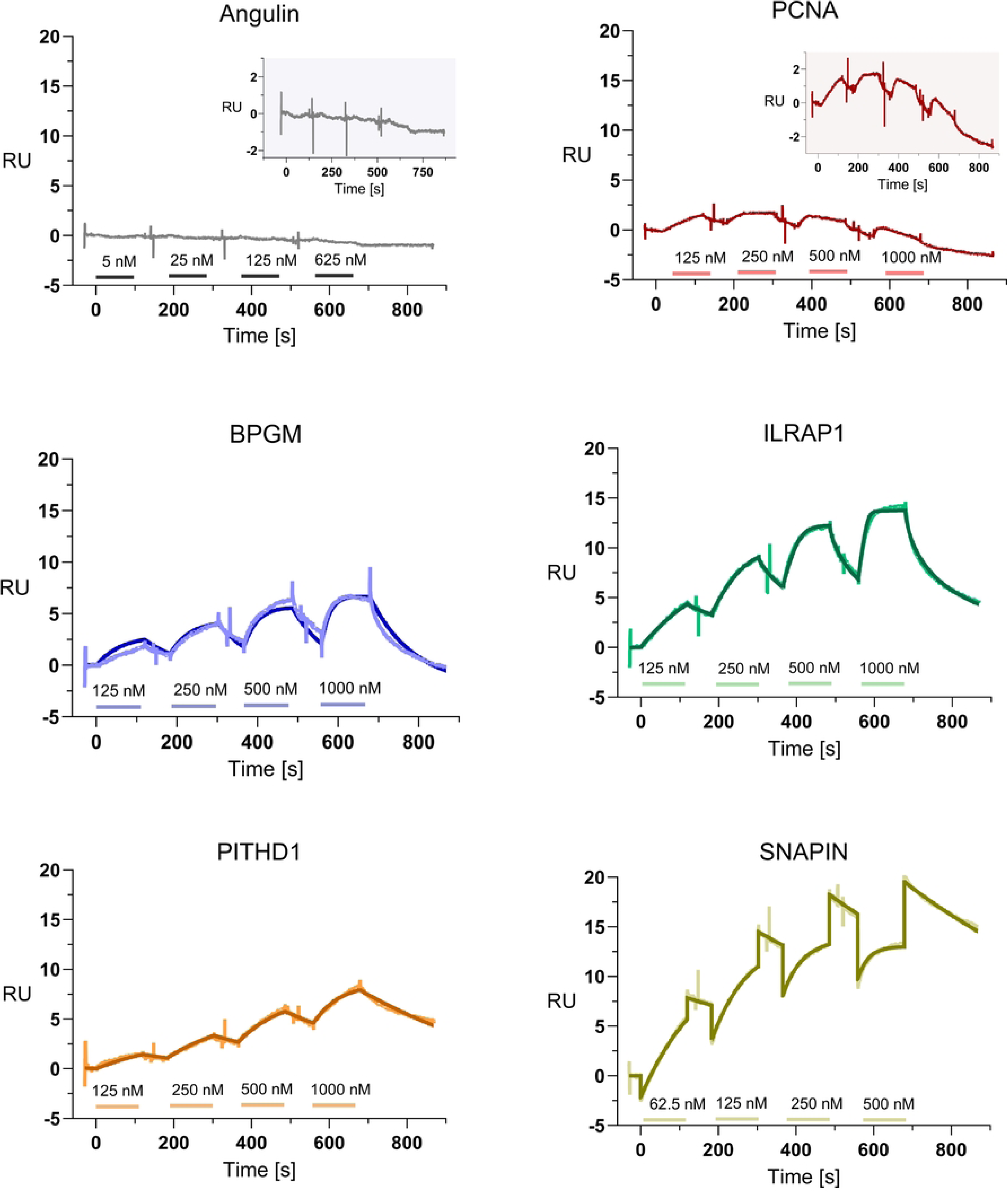
Interaction analysis of TiPs to human 2N4R tau by SPR. BPGM, ILRAP1, PITHD1, and SNAPIN showed concentration-dependent binding to human 2N4R tau, whereas PCNA exhibited no concentration-dependent behavior and consistently low binding levels, indicating a lack of interaction. Angulin 1, which was used as a negative control, also showed no interaction. Analytes were injected at increasing concentrations as indicated. Horizontal lines mark the injection phases. Experimental measurements and fits (fitted with a 1:1 binding model) are presented as lighter pointed and darker solid lines, respectively. Responses of Angulin 1 and SNAPIN are highlighted in insets.

**Table 4.** Kinetic parameters of the interaction of selected TiPs with human 2N4R tau.

| **Analyte** | **k<sub>a</sub> (1/Ms)** | **k<sub>d</sub> (1/s)** | **K<sub>D</sub> (M)** | **Rmax (RU)** |
|---|---|---|---|---|
| PITHD1 | 1.16 × 10<sup>4</sup> | 2.06 × 10<sup>-3</sup> | 1.77 × 10<sup>-7</sup> | 12.3 |
| SNAPIN | 7.21 × 10<sup>4</sup> | 1.56 × 10<sup>-3</sup> | 2.16 × 10<sup>-8</sup> | 20.4 |
| BPGM | 3.83 × 10<sup>4</sup> | 9.76 × 10<sup>-3</sup> | 2.55 × 10<sup>-7</sup> | 10.1 |
| ILRAP1 | 6.16 × 10<sup>5</sup> | 8.97 × 10<sup>-2</sup> | 1.46 × 10<sup>-7</sup> | 14.8 |

Notes: The association (k<sub>a</sub>), dissociation rate (k<sub>d</sub>), and equilibrium dissociation K<sub>D</sub> constant were calculated by performing a 1:1 binding fit on sensograms. Rmax - maximum theoretical response, referring to the binding capacity.

## Discussion

The primary objective of this study was to identify novel tau interaction partners using the yeast two-hybrid system. Across four independent screens - employing human full-length tau (2N4R) alongside three of its fragments (N-terminus, MTBD, and C-terminus) - a total of 418 unique clones were initially isolated. From the primary screen of 418 clones, only 58 clones were subsequently validated as true positives, highlighting the necessity of high-stringency selection steps. Several high-throughput Y2H screening workflows developed in past studies omitted stringent validation phases, thereby leading to elevated false-positive rates [93, 94]. Following the exclusion of duplicates, non-coding fragments, uncharacterized proteins, pseudogenes, truncated peptides, and out-of-frame fusions, we identified 15 novel tau-interacting partners (Table 1).

The screening utilizing the tau C-terminus had the highest hit rate, identifying eight novel interactors. The remaining identified TiPs were distributed across the screens as follows: two bound to full-length 2N4R tau, four to the N-terminus, and three to the MTBD (Table 2). ATP1B1 was concurrently identified as a binding partner for both 2N4R tau and the C-terminus. Screening with the pathological tau fragment (human Tau151–391) yielded no viable results. Following repeated screening attempts, post-mating cell death occurred, highly likely due to the fragment’s cytotoxicity, which led to the termination of the screen.

The high false-positive rate (94–96%) and the high frequency of duplicates can be attributed to the architecture of the cDNA library, which contains short, fragmented cDNA inserts that encode short peptides rather than intact, full-length proteins [95]. Furthermore, post-mortem RNA degradation and the inherent presence of non-coding regions or pseudogenes in cDNA libraries could increase false-positive rates, ultimately limiting the yield of authentic novel interactors.

For most of the discovered TiPs, the interacting sequences were well conserved across multiple protein isoforms. A notable exception was IL1RAP, which interacted exclusively with the soluble cytoplasmic isoform (sIL1RAP) rather than with the membrane-bound isoform. Two of our identified proteins overlapped with those from a previous C-terminal tau Y2H screening conducted by Lee et al. using the same cDNA library: PCNA and SNAPIN [15]. We also captured PCNA with the tau C-terminus, but SNAPIN was identified in our study using the full-length tau bait. In our previous experiments, restricted to tau N-terminus Y2H screening using the same library, we isolated only a single partner, CHORDC1 [24], in contrast to the study by Lee et al., in which it was identified with the tau C-terminus [15].

In our earlier report, we identified tau-interacting proteins in the brains of the SHR72 rat model of tauopathy using an *in vivo* cross-linking approach coupled with immunoprecipitation-mass spectrometry (IP-MS) [16]. In both the previous and current screening approaches, only one protein, ATP1B1, emerged as a shared hit without validation. All 15 proteins identified in this study are considered novel tau interaction partners, as they are not listed in the BioGRID interaction database for validated low-throughput experiments [12]. Although CACYBP is already present in the BioGRID database, its inclusion resulted from a high-throughput experiment lacking further validation [96].

Our screening did not discover any previously established tau interactors. This absence can be attributed to several standard drawbacks of the Y2H system, including protein misfolding, steric hindrance of the reconstituted transcription factor, the absence of some mammalian post-translational modifications (PTMs) in yeast, a requirement for mammalian-specific co-factors, or restricted nuclear translocation. Furthermore, severe target cytotoxicity or autoactivation can lead to yeast cell death, selectively eliminating certain interactors [97]. The high stringency of our screening may also filter out genuine but transient interactions, resulting in false negatives [98]. Additionally, post-mortem mRNA degradation and subsequent cDNA library normalization could lead to a loss of total transcript diversity [99, 100].

Orthogonal validation of the 15 novel TiPs was essential to contextualize the yeast two-hybrid (Y2H) screening data within true mammalian physiology, accounting for the artificial overexpression and distinct post-translational modifications inherent to yeast models. A prime example of this physiological relevance is the identified interaction between the N-terminus of tau and the extracellular domains of proteins such as ATP1B3. While tau is traditionally defined as an intracellular protein, its low-concentration presence in the extracellular space is increasingly recognized as playing a role in cell-to-cell signaling and the trans-synaptic spreading of tau pathology [101].

Our multi-platform validation strategy revealed that these interactors are not static bystanders but dynamic participants that behave uniquely based on species, tissue type, and disease state [102]. For instance, the persistent 100 kDa band observed for IL1RAP across both human and rat cohorts—far exceeding its expected 42 kDa soluble form—strongly implies that tau could preferentially bind its membrane-associated isoform, suggesting a potential role for this interaction beyond a purely soluble neuroinflammatory cascade toward a membrane-bound receptor pathway.

Our observed discrepancies between physiological and pathological conditions reveal a molecular separation between homeostatic preservation and neurodegenerative progression. The selective presence of ATP1B3 and NCAM1 in healthy control brains and their complete absence in AD tissues point to a possible disease-driven loss of function that could exacerbate disease progression. The different conformational preferences of individual TiPs highlight their distinctive roles as neuroprotectors or participants in neurodegeneration. Proteins like ATP1B1 play fundamental roles in tau biology, as evidenced by their cross-species validation and persistent colocalization even at basal expression levels in human neuroblastoma cell lines. Interactors such as PCNA and PDE4DIP selectively target fragmented, pathological tau across neuroblastoma cell lines. This selective binding implies that they are actively recruited to or driven by the structural alterations that characterize hyperphosphorylated or truncated tau species.

The various outcomes across different validation platforms underscore the transient, context-dependent nature of the tau interactome, rather than experimental failure. Interactors that failed in complex tissue co-immunoprecipitations but succeeded in high-sensitivity neuroblastoma cell models, such as PCNA and PDE4DIP, highlight the technical challenges of capturing low-abundance or highly localized signals in post-mortem tissue that may be subject to proteolysis [103]. For natively nuclear proteins like PCNA, which maintains a low abundance baseline in post-mitotic neurons, is this interaction likely upregulated only during acute nuclear stress, such as DNA damage or pathological cell-cycle reentry [76]. Ultimately, these differential validation patterns prove that tau’s macromolecular landscape changes dynamically, resulting in possible shifts of physiological interactions in cellular homeostasis to active participation in pathological neurodegeneration.

Biophysical characterization via SPR successfully transformed yeast two-hybrid screening data into quantified molecular kinetics, confirming direct physical affinities for a targeted subgroup of novel TiPs. The successful purification and kinetic description of SNAPIN, BPGM, PITHD1, and IL1RAP provide robust biochemical evidence that tau directly engages with proteins that control vital cellular pathways, including vesicular trafficking, systemic metabolism, and neuroinflammation. Among these, the novel interactor SNAPIN formed the most stable complex with full-length tau, exhibiting the highest overall binding affinity. Under healthy homeostatic conditions, this tau-SNAPIN complex is likely to participate in physiological endolysosomal trafficking and synaptic vesicle recycling. In the AD brain, hyperphosphorylation or aggregation of tau could physically disrupt this high-affinity interaction, leading to endosomal-autophagic dysfunction and synaptic loss characteristic of advanced neurodegeneration [104].

The negative outcome for PCNA in SPR experiments, which mirrors its absence in tissue co-immunoprecipitations, provides important clues about its conditional molecular role. Rather than dismissing PCNA as a screening artifact, its inability to bind full-length tau *in vitro* suggests that the interaction is most likely conditionally-driven. In a healthy, mature brain, neurons are post-mitotic, and both PCNA abundance and its interaction with tau are unlikely to be present. However, in the AD pathological environment, chronic oxidative stress triggers abortive cell-cycle reactivation and severe nuclear stress. We hypothesize that under these active pathological conditions, PCNA is mobilized and transiently complexed with specific altered tau conformers to modulate DNA repair pathways or cell survival, shortly before apoptosis occurs. Thus, the tau-PCNA pathway may represent a latent feedback loop in pathological conditions, which is completely absent in healthy tissue or is simplified in in *vitro* setups.

Expanding these findings through *in silico* network modeling of our cumulative set of 21 interactors reveals a vast macromolecular landscape with over 6,900 interactions, including a critical sub-network of 342 nodes directly linked to AD pathogenesis. Crucially, the identification of primary nodes like IL1RAP, NCAM1, NPTX1, and PCNA—which possess established, multi-pathway links to AD within known molecular cascades—demonstrates that tau’s interactome is deeply intersected with systemic neurodegenerative pathways. In the healthy brain, these proteins operate within balanced networks to maintain synaptic plasticity, resolve inflammation, and preserve genomic stability. In AD, tau pathology acts as a molecular keystone, disrupting these interconnected networks simultaneously. By mapping these high-affinity biophysical bindings to a larger disease-associated interactome, these findings highlight how localized tau misfolding can propagate outwards, driving the systemic network failure that characterizes the transition from healthy cognitive aging to clinical Alzheimer’s disease.

In the context of therapeutic approaches for neurodegeneration, four proteins stand out as possible targets – SNAPIN, CACYBP, sIL1RAP and ATP1B3. SNAPIN is essential for retrograde endosomal-lysosomal transport and vesicle release, whereas axonal autophagic stress is a major driver of neurodegeneration. Upregulation of SNAPIN could accelerate the removal of misfolded proteins and drive the delivery and degradation of toxic tau aggregates in the lysosome. The CACYBP/SIP complex plays a role in neuroprotection due to its dephosphorylating activity, which is suppressed by other Ca^2+^-binding proteins (e.g., S100A6). Inhibiting the interaction between CACYBP and S100A6 could increase CACYBP/SIP dephosphorylation activity and, in turn, its neuroprotective role. Because sIL1RAP and ATP1B3 are present in the extracellular space, we suggest that their interaction with pathological tau species could inhibit disease propagation by blocking pathological tau seeding.

The most critical discovery of this study is the identification and multi-platform validation of an expanded, multi-compartment tau interactome, highlighted by the discovery of 15 novel interaction partners spanning the nucleus, cytoplasm, and extracellular space. By bridging genetic screens (Y2H), tissue-specific biochemistry (co-IP), and real-time biophysics (SPR). Our findings contribute to the growing consensus that tau pathology does not operate in isolation. Instead, they provide further evidence that tau physically engages with, and likely disrupts, vital cellular networks governing proteostasis, synaptic trafficking, and metabolic homeostasis. Many of these pathways are heavily compromised in Alzheimer’s disease. These physical interactions with tau likely contribute to downstream mitochondrial decay, impaired autophagy, disrupted membrane potential, and non-canonical tau secretion. Furthermore, these findings link tau to altered metabolic pathways and cellular scaffolding networks, providing a useful framework for future targeted functional studies in tauopathies.

### Limitations of the Study

While this study provides a comprehensive workflow for the discovery and biophysical validation of novel tau interaction partners, several technical limitations must be acknowledged.

First, although *Saccharomyces cerevisiae* is an effective eukaryotic model for Y2H screening, its endogenous post-translational modification machinery differs significantly from that of mammals, particularly humans. Furthermore, because the commercial cDNA library utilized in this work contained fragmented cDNA transcripts rather than full-length open reading frames, some interactions may be restricted to isolated domains or brief peptide fragments. The classical transcription-based Y2H system is also inherently constrained when evaluating membrane-bound proteins or receptors. Because the system forces interactions to occur within the yeast nucleus, it can occasionally yield false-positive configurations that are physiologically impossible *in situ* due to divergent cellular compartmentalization.

Second, the *in vitro* orthogonal validation strategies carry platform-specific constraints. Immunofluorescence colocalization assays are limited by the intracellular environment, making it technically infeasible to capture extracellular interactions or secreted protein domains.

Third, regarding biophysical quantification via SPR, the specific PTM profiles imparted on the recombinant proteins by the CHO secretory system remain uncharacterized and may deviate from endogenous human modifications. Additionally, the presence of minor trace contaminants or non-specific proteins within the purified fractions could introduce subtle parasitic binding or alter sensorgram kinetics.

Finally, this study was designed strictly to establish direct physical contact and kinetic parameters, without defining downstream biological functions. Our findings are then primarily descriptive and characterizing, as functional knock-out models and *in vivo* phenotypic assays are absent. Future functional studies are warranted to elucidate the precise pathological and physiological consequences of these novel complexes.

## Conclusions

Our study maps an expanded tau interactome by identifying 15 novel physical interactors across diverse cellular pathways. By establishing tight biophysical links to critical modulators such as SNAPIN and CACYBP, as well as to extracellular targets, these findings provide a mechanistic framework for understanding how tau may drive proteostatic, synaptic, and metabolic decay. Translating these interactions into targeted therapeutics—such as accelerating autophagic clearance, inducing targeted tau dephosphorylation, or deploying extracellular decoy receptors—offers a promising frontier to halt disease progression. Ultimately, these newly uncovered partnerships provide valuable biochemical foundations to guide future functional characterization and drug discovery efforts in Alzheimer’s disease and related tauopathies.

## Acknowledgments

The authors would like to thank Mgr. Silvia Bagelova Polakova, PhD from the Institute of Animal Biochemistry and Genetics, SAS, Bratislava, for her valuable assistance and support throughout Y2H screening procedures.

---

**Fig. S1.**
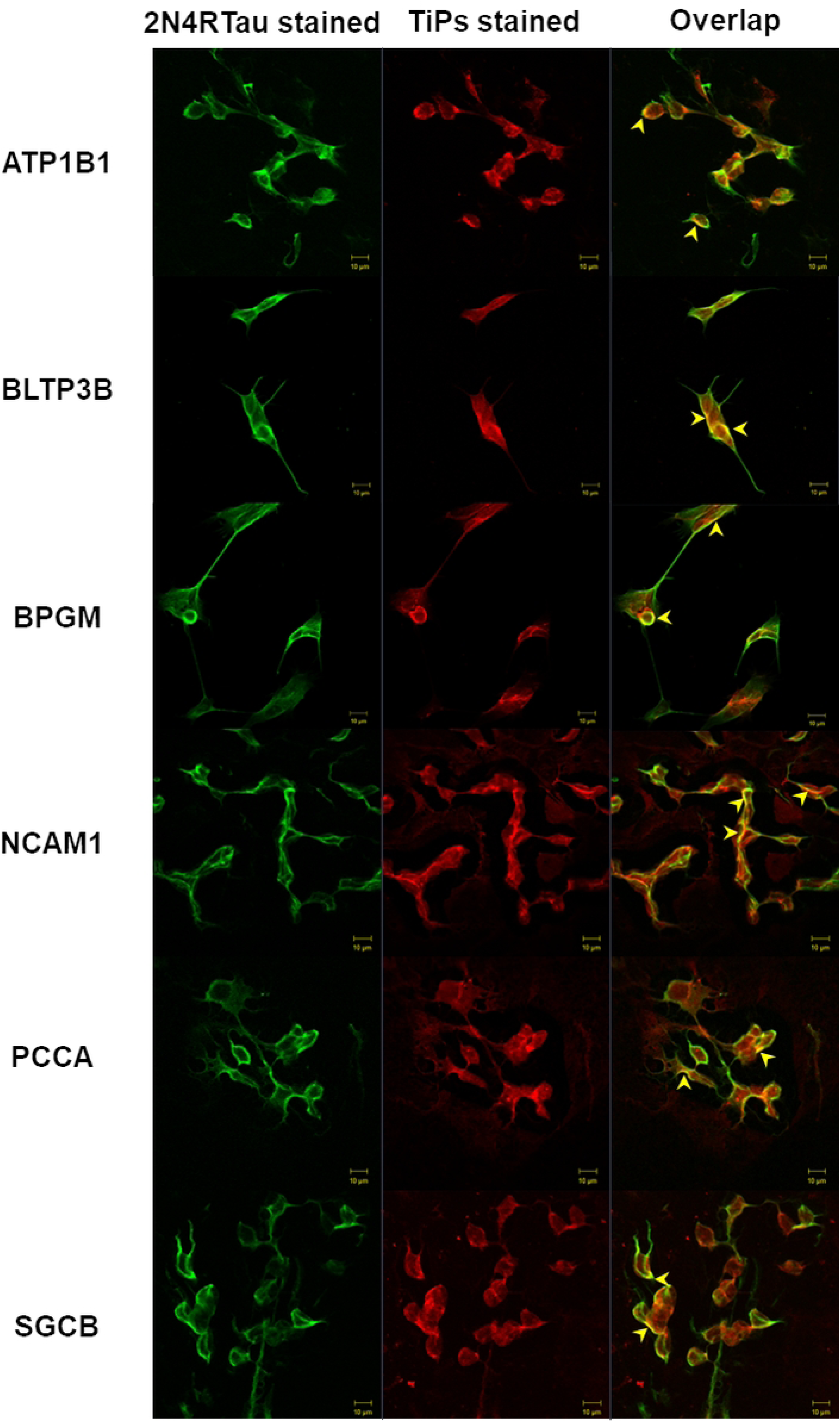

**Fig. S2.**
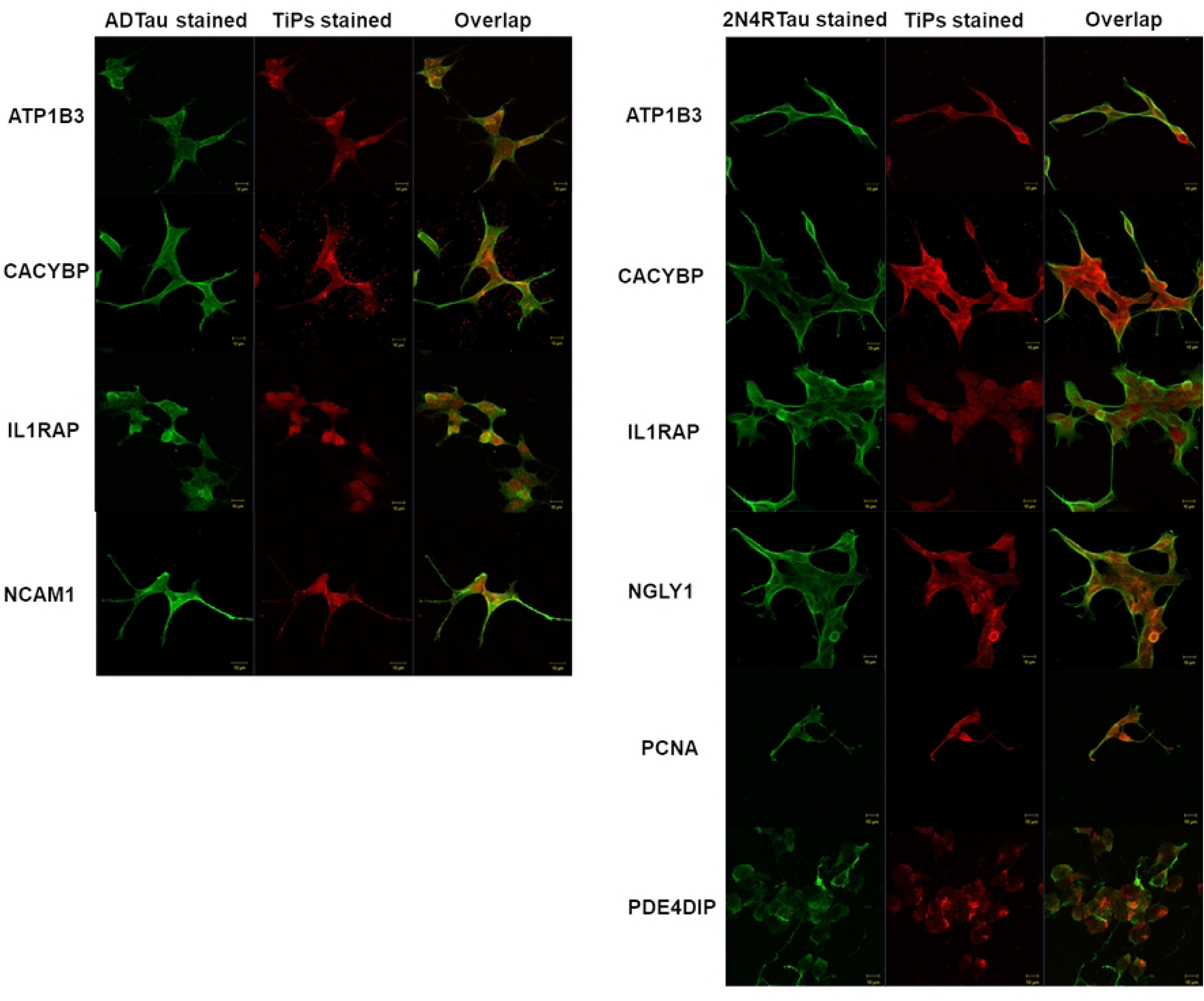

**Fig. S3.**
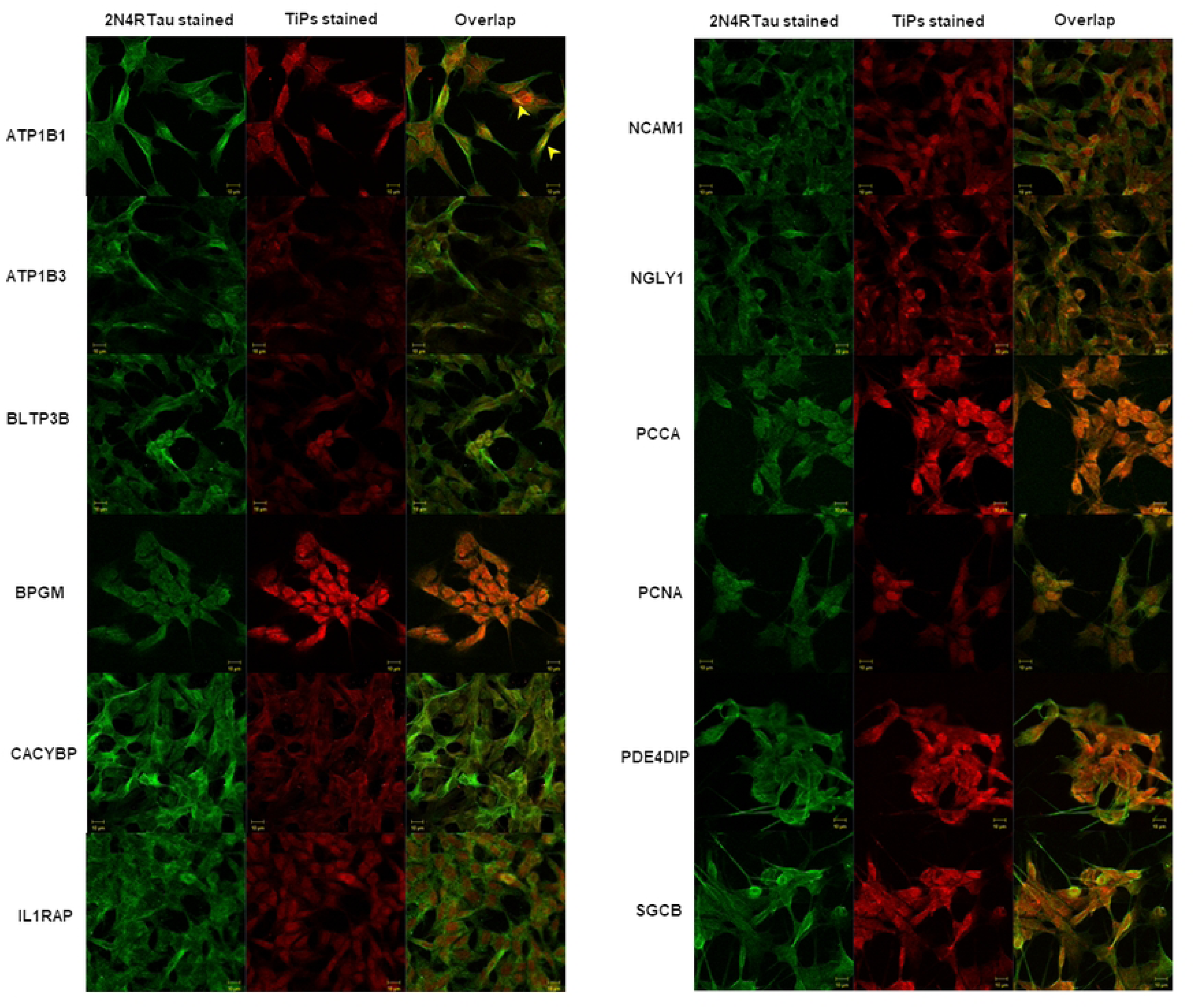

